# A capture assay to analyse immunoglobulin secretion at the single-cell level

**DOI:** 10.64898/2026.01.07.698251

**Authors:** Theresa H. Ward, Grégoire Altan-Bonnet

## Abstract

Several secretion capture assays have been developed to measure a secreted protein of interest, using paired monoclonal antibodies of different specificities that bind simultaneously to a cell surface antigen and to the secreted protein of interest. Here we have developed a novel method to measure antibody secretion on B cells, which, by capitalising on the bivalent nature of an antibody molecule, utilises an anti-immunoglobulin (Ig) antibody to bind on one side to the membrane-bound Ig (in the B cell receptor), and then capture any secreted Ig with the second binding site. Binding of the secreted cargo is then detected through a second application of anti-Ig, this time fluorophore-labelled. We demonstrate that this can effectively measure single cell Ig secretion in a population of B cells undergoing differentiation to plasma cells. This offers a reliable and accessible tool for quantifying antibody secretion dynamics at the single-cell level with potential applications in immunological studies, antibody discovery and functional profiling of B cell responses.

## Introduction

Plasma cells are terminally differentiated B lymphocytes that are highly specialised to synthesise and secrete large amounts of immunoglobulins^1^. These immunoglobulins, or antibodies, provide humoral immunity to combat infectious diseases. They are also an important tool in immunotherapeutics. Robust and accurate secretion controls are required to ensure that the antibody molecules are correctly assembled, modified and released from cells.

Identification of high-secretion human clones is often a time-consuming process. Currently immunohistochemistry and flow cytometry are the two main techniques allowing the identification of secreting cells^2,3^, but generally this entails a potential block in the secretory pathway to accumulate secreted protein inside the cells as a measure of how much is being produced, and then fixation/permeabilization of the cells in order to detect the intracellular accumulation. Alternatively, a secretion-capture assay, similar to the ones used to detect cytokine secretion^4-6^, would enable the measurement of secretion from living cells. Some live secretion assays require capture of the secreted protein within a matrix while others need development of a cell receptor-capture hybrid antibody^7-9^. Here we have developed a simple direct live-cell capture assay to enable identification of high level IgM-secreting plasma cells in an efficient, easy and cost-effective method, using readily available reagents. Rather than using a hybrid conjugate antibody to tether the capture antibody to the cell, we make use of the multivalent binding capability of the capture antibody, using a polyclonal mixture of anti-IgM antibodies, to tether the antibody to the cell surface B cell receptor and then capture secreted IgM as it is released. We envisage that this might enable several downstream applications that could be applied to other immunoglobulin isotypes including (i) identification of secretion in germinal centre developing B cells; (ii) a tool to read out the effect of investigations into B cell protein function on antibody assembly and secretion and (iii) enabling high throughput screening and isolation of Ig-producing cells to facilitate development of high-producing cell lines.

## Methods

### Isolation and stimulation of peripheral human B cells

Fresh, anonymous, human donor blood was obtained under ethical consent from healthy volunteers either through the NIH blood bank or through the donor scheme at LSHTM (ethics reference 17854). Peripheral Blood Mononuclear Cells (PBMC) were isolated by density centrifugation over Ficoll Histopaque-1077. CD20^+^ B cells were positively selected using MACS Microbeads (Miltenyi Biotec, Auburn CA), according to the manufacturer’s instructions, and counted. Immediately upon isolation, B cells were diluted to 1 x 10^6^ cells/ml in B cell medium (HEPES-buffered RPMI, 10% FCS, 2 mM glutamine, 0.1 mg/ml streptomycin, 100 U/ml penicillin), supplemented with the combined antigen mixture of 100 µg/ml SAC, 10 µg/ml PWM, and 1 ng/ml CpG2006 to induce differentiation^10^, and incubated at 37°C with 5% CO_2_. Cells were analyzed by removal of an appropriate aliquot from the stimulated culture at the timepoints indicated in the text – over a period of up to 6 days – and processed according to the experimental method.

### IgM capture assay

Cells were collected from the *in vitro* culture, washed in assay buffer (PBS, 1% FCS), incubated on ice with 50 µg/ml capture antibody in 50 μl assay buffer for 15 min, washed and re-suspended in 150 µl culture medium and incubated for times as shown. Cells were then collected, washed, and stained for IgM secretion using the detection antibody at 5 µg/ml together with either DAPI or Live-Dead stain. Dead cells were excluded by DAPI inclusion or Live-Dead stain (Invitrogen, Carlsbad CA) and fluorescence was analyzed by flow cytometry.

### Bead ELISA

We developed a simple bead-based assay to quantify the amount of IgM secreted in the supernatant of cells. 10 µm diameter latex beads (Bangs Laboratories, Fishers IN) were washed in PBS, then coated with a polyclonal solution of anti-IgM antibodies (Jackson Immunochemicals, West Grove PA) for 30 min at room temperature. The beads were then spun and washed twice with complete medium and resuspended at 10^5^ beads/ml. 1000 beads were added with the supernatant under consideration and incubated for 30 min on ice. The same procedure was repeated with a solution of human IgM to serve as a calibration curve. Beads were then spun, washed, then resuspended in FACS buffer (4% fetal calf serum in PBS + 0.1% w/v sodium azide). A solution of fluorescently-labeled anti-IgM was then added to the beads and incubated for 30 min on ice. Beads were then again spun, washed, then resuspended in FACS buffer.

### Flow cytometry

Acquisition was performed on a BD LSRFortessa™ Cell Analyzer (BD Biosciences, San Jose CA) or Cytek^→^ Aurora (Cytek, Bethesda MD). Data were analyzed using FlowJo™ software (*Version 10*.*1*, Tree Star, Ashland, OR) and Plateypus, a custom Python pipeline^11^.

### Antibodies and reagents

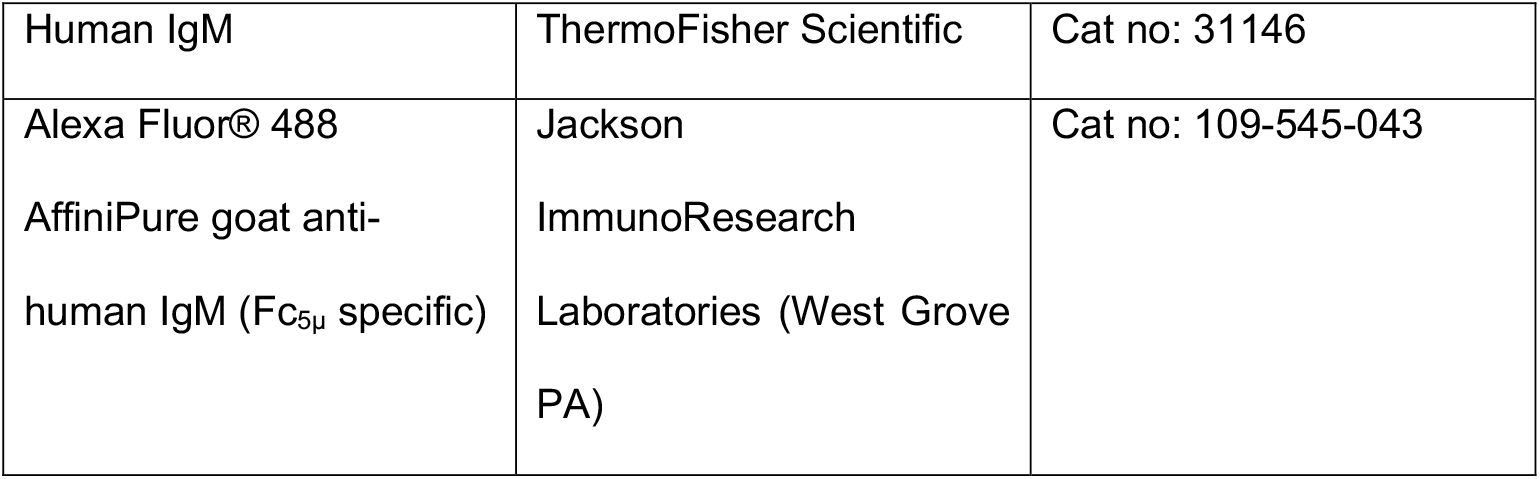
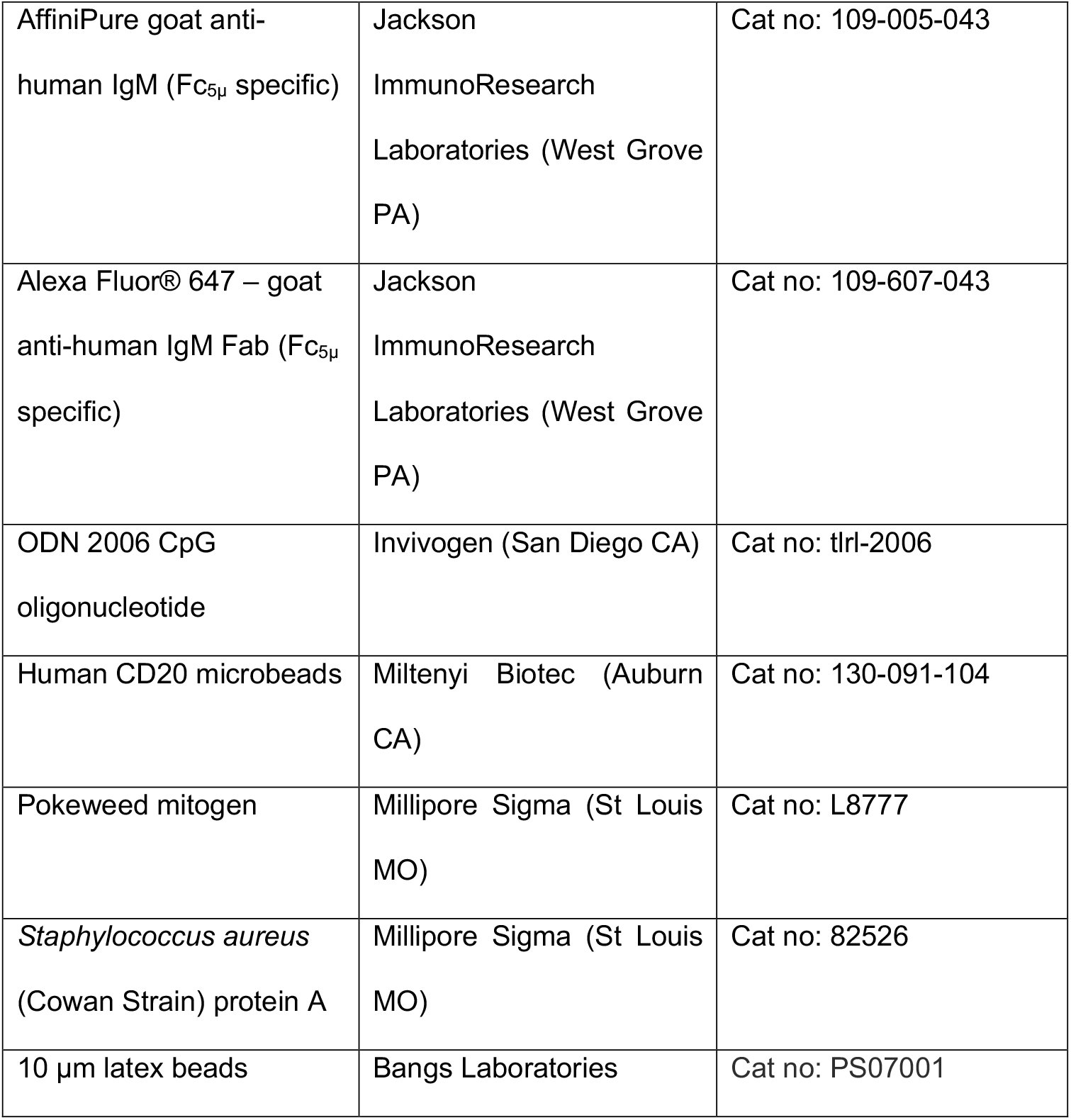

## Results

Several secretion capture assays have been developed to measure a secreted protein of interest using paired monoclonal antibodies of different specificities that bind simultaneously to a cell surface antigen and to the secreted protein of interest. These antibodies can be paired either through direct covalent coupling or through co-coupling to nanoparticles. The cell surface antigen must be present on the cell of interest and would be minimally reactive to binding of the coupled antibody.

To measure antibody secretion on B cells, our approach, rather than using two antibodies – one for binding to the cell and one for capture, was to use the property of antibody-producing cells to synthesize the same immunoglobulin in both membrane-bound (mIg) and secreted form (sIg) differing only at the carboxyl terminus^12^. The membrane-bound Ig molecules form part of the B cell receptor expressed on the cell surface. Differential processing of the primary Ig transcript determines whether a secreted or membrane-bound antibody will be produced but the large part of the antibody molecule remains the same. Therefore an antibody that would bind to mIg on the cell surface would also bind to any secreted antibody. By capitalising on the bivalent nature of an antibody monomer, the anti-Ig antibody would therefore be able to bind on one side to the membrane-bound Ig (in the BCR), and then capture any secreted Ig with the second binding site. Binding of the secreted cargo is then detected through a second application of anti-Ig, this time fluorophore labelled.

The ability of the anti-IgM to effectively capture IgM was first tested in the Burkitt-lymphoma-derived Ramos B cell line that natively expresses both membrane-bound and secreted IgM antibodies^13^. AlexaFluor(AF)488-tagged anti-hIgM was added to the Ramos cells to demonstrate labelling of surface mIgM (Fig. 1A). To determine whether IgM might be captured with the bivalent anti-IgM, Ramos cells were first incubated with anti-hIgM (the capture step), then exogenous hIgM was added as a binding substrate, and finally AF488-tagged anti-hIgM added to detect captured sIgM. As a control, no substrate IgM was added (Fig. 1B). The control checks that there is no binding of AF488-anti-hIgM to the cells in the absence of substrate capture. Concentration of the capture antibody stage was tested through titration of anti-hIgM (Fig. 1C) with concentration of 50µg/ml determined to be effective.

**Figure 1.**
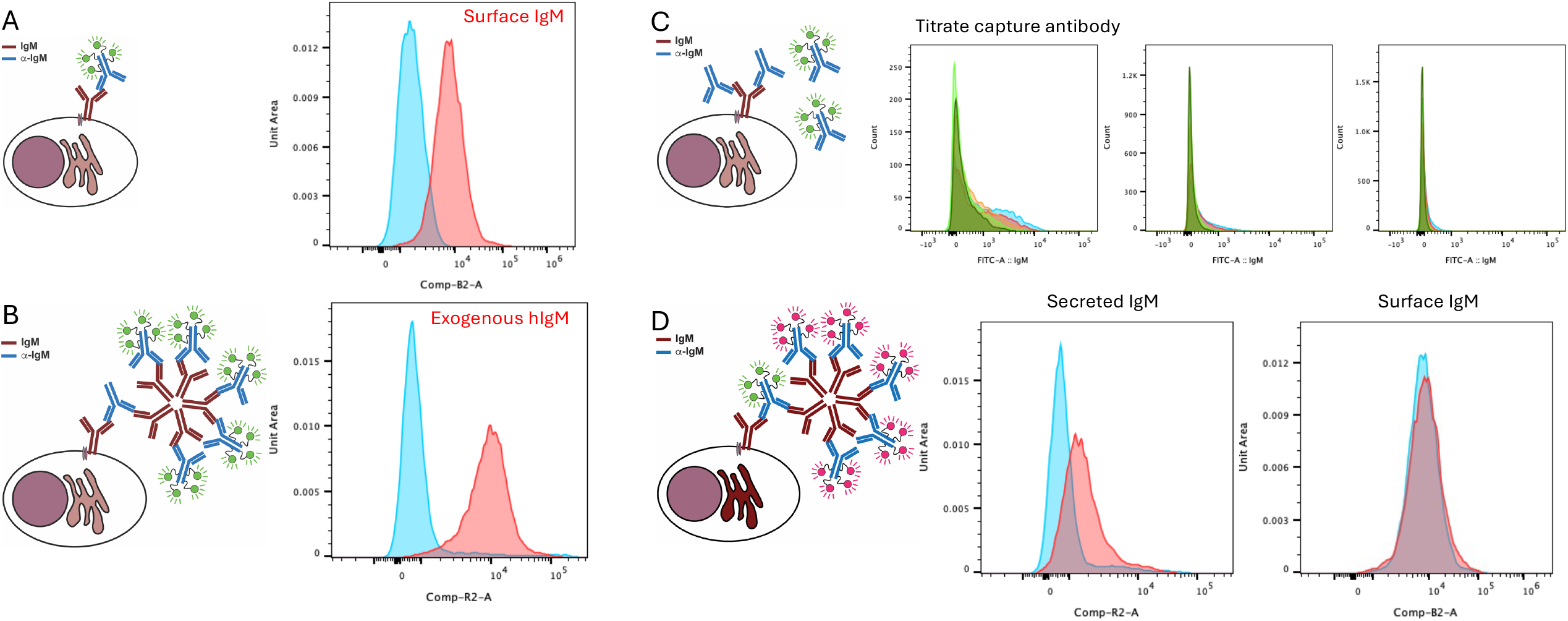
A functional capture assay on Ramos cells. Cartoons are used to demonstrate the IgM molecules produced by the cells of interest and the anti-IgM molecules applied as capture and detection molecules in the assay. A. Detection of surface IgM with anti-IgM. Blue, control no fluorescently antibody applied, red AF488-anti-IgM added to cells. B. Cells were incubated with unlabelled capture anti-IgM, then exogenous human IgM was added to the cells as a control to bind to the capture antibody; binding was then detected with AF647-anti-IgM. C. Titration of capture IgM. Increasing quantities of capture antibody (unlabelled anti-IgM) were titrated against AF488-anti-IgM until only background AF488-anti-IgM binding was detected (left to right 1 µg/ml, 10 µg/ml, 50 µg/ml). D. The dual color assay uses a labeled capture antibody and a second labelled detection antibody. Ramos cells incubated either on ice (blue) or at 37°C (red) for 2 hours, following incubation with the capture antibody, were then incubated with detection antibody and analysed by flow cytometry.

Capture of endogenous IgM secretion was tested through first binding anti-hIgM to surface BCR in Ramos cells and then incubating either on ice or at 37°C for 2 hours (Fig. 1D). This showed that upon incubation on ice, secretion was inhibited while there was a distinctive shift in fluorescence in the cells incubated at 37°C to demonstrate secretion of IgM over the time period. To evaluate whether this might be related to turnover of surface IgM, staining of the surface BCR was incorporated into the assay using a dual color protocol, with the blocking antibody labelled with AF488 and the detection antibody labelled with AF647. No change in the mIgM was found during the time course demonstrating that the capture was related to the secretion of IgM.

We next tested the assay in primary human B cells. Purified CD20^+^ peripheral primary human B cells were cultured with a mitogenic antigen mixture for 6 days^10^. At the indicated timepoints, cells were removed and IgM secretion assessed using the capture/detection assay. In Fig. 2A, the unstimulated B cells at day 0 show no secretion, the profile of the cells incubated at 37°C matches the cells on ice (negative control), while those incubated with exogenous hIgM show that the cells carry functional B cell receptor that is able to capture IgM (positive control). Following 3 days of stimulation, some cells show low levels of secretion, while by day 6 this becomes enhanced, with clearly increased numbers of highly-secreting cells. There is also a clear increase in capture between cells incubated for 1 hour at 37°C versus those incubated for 4 hours at 37°C, therefore increasing time of assay increases secretion detection. This becomes much closer to the maximal secretion that is measurable using the exogenous hIgM loading control. Incubation on ice inhibits the secretion process. Fig. 2B demonstrates that the AF488-labelled mIgM is not altered between ice- and 37°C-incubated cells at different timepoints while the AF647-labelled sIgM distinctly changes, therefore the increase of signal is due to secretion rather than cycling of the capture antibody-labelled mIgM.

**Figure 2.**
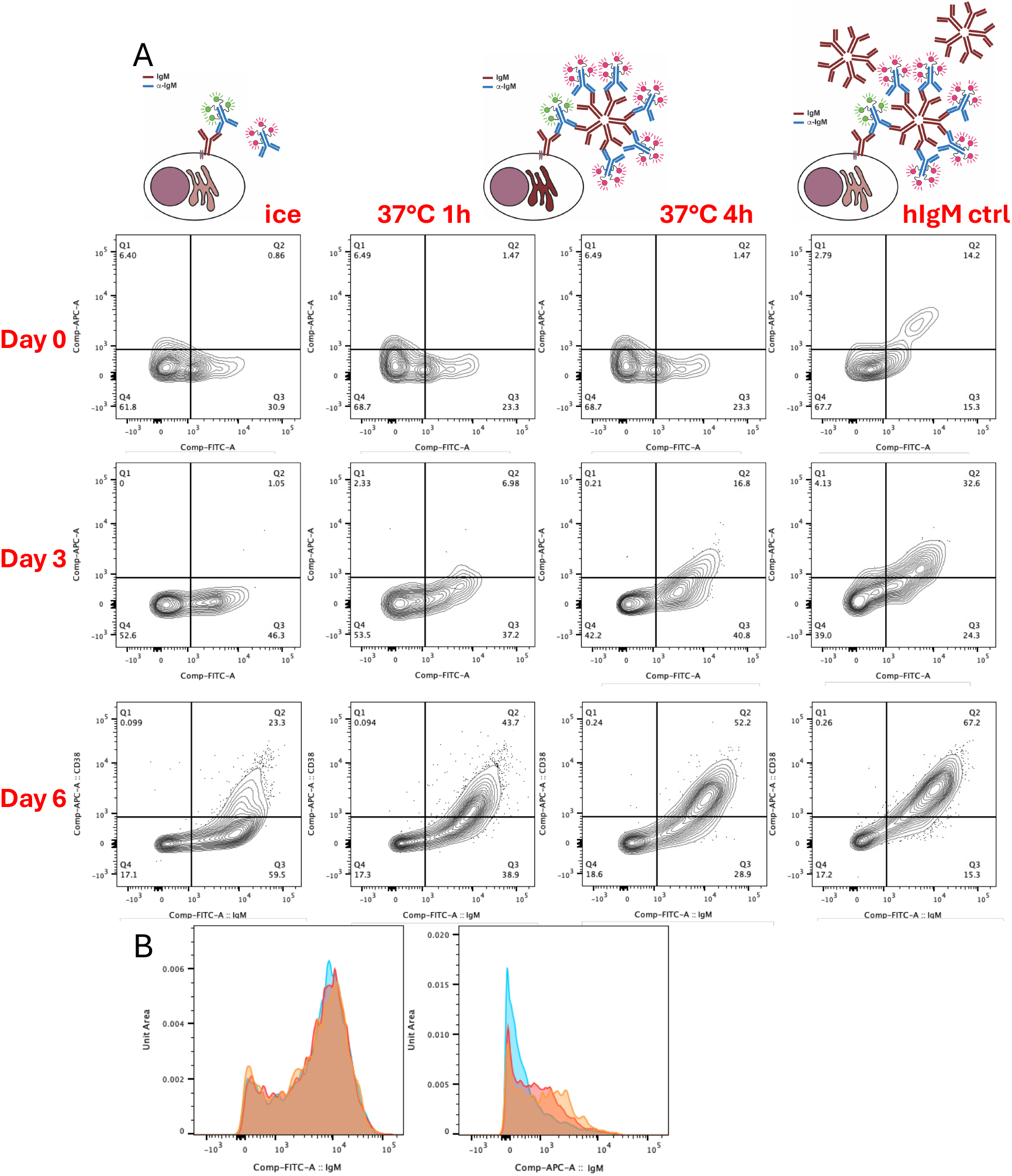
Capture assay on primary human B cells. Utilising the blocking and capture antibodies with different fluorophores, primary human B cells were purified (day 0) and then activated with a mitogen mixture for the given time periods. A. Secretion was tested on cells from different time points and cells were either incubated on ice (no secretion control), at 37°C for 1 hour or for 4 hours, and with addition of exogenous hIgM (positive control). B. The left panel shows AF488-anti-IgM labelling of mIgM; right panel shows AF647-anti-IgM detection of sIgM (blue ice, red 1h, orange 4h, d6 cells).

To demonstrate that the capture is dependent on secretion from the cells, we used brefeldin A to block trafficking from the secretory pathway and therefore Ig secretion. In Fig. 3A, IgM capture during BFA treatment closely compares to cells incubated on ice at all timepoints, while cells incubated at 37°C without a secretory block show increased IgM capture, demonstrating that BFA significantly reduces captured sIgM. In Fig. 3B, detection of IgM in permeabilised cells shows a significantly increased internal pool only in day 6 cells, which would reflect the IgM trapped by BFA treatment, and hence lack of captured sIgM in the live cell assay. If BFA treatment might result in less delivery of mIgM to the cell surface, then this might result in less capture of sIgM. In Fig. 3C, capture antibody applied to the cells before or after incubation at 37°C ± BFA remains the same. This demonstrates that there is no change in levels of surface mIgM during the timecourse, including upon treatment with BFA.

**Figure 3.**
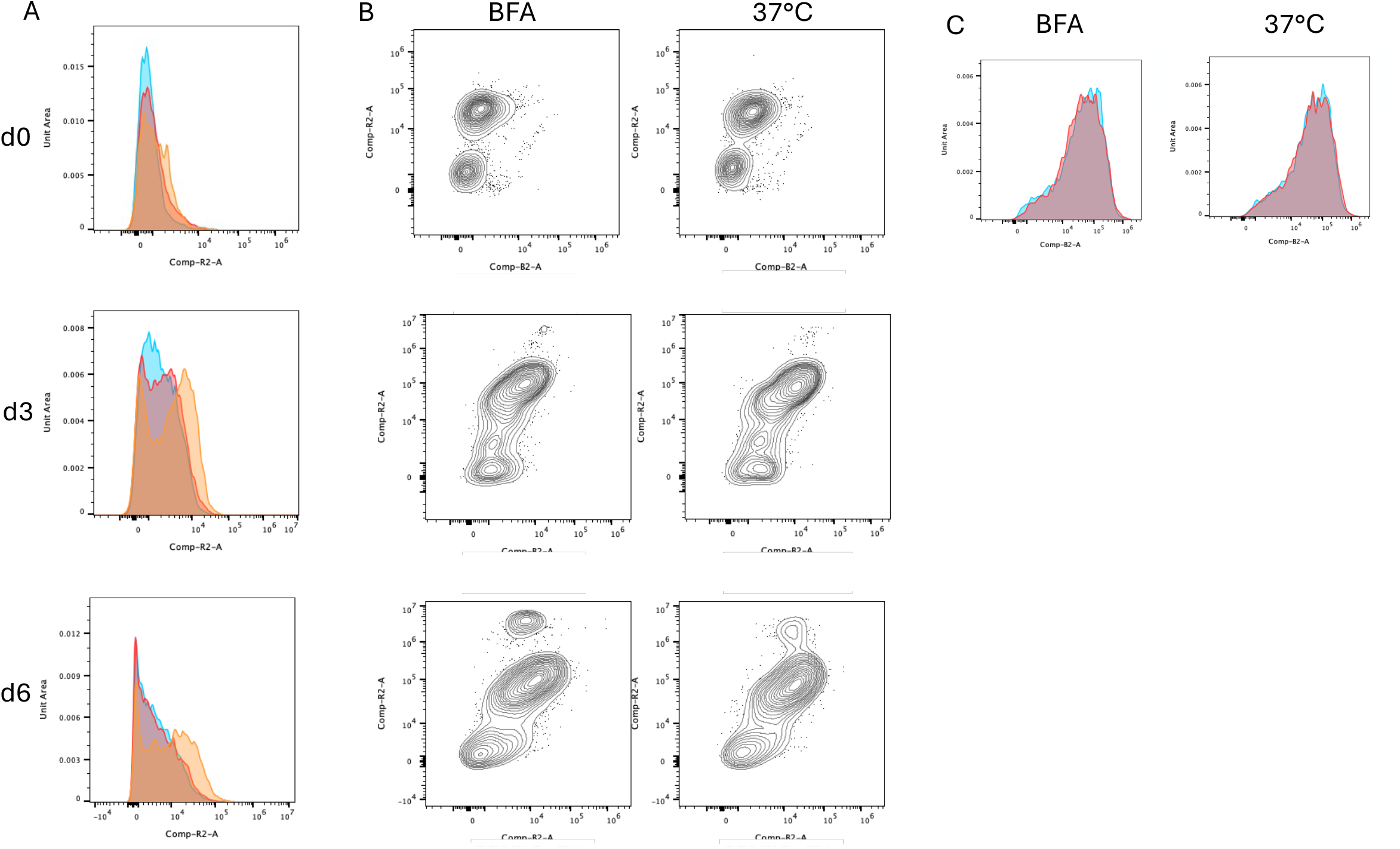
Inhibition of secretion with brefeldin A. A. B cells at differentiation time points as shown underwent the secretion assay with incubation steps of 2hours. blue ice, red BFA at 37°C, orange 37°C. B. endogenous IgM detected following fixation, permeabilisation and staining. C. mIgM was labelled with with AF488-anti-IgM before or after incubation at 37°C ± BFA, showing no change over the time course (blue labelling before incubation, red labelling after incubation).

To compare on-cell capture of IgM to supernatant measurements, we utilised a bead assay to quantify IgM concentration in cell supernatants collected at given timepoints. Fig. 4A demonstrates increase in IgM concentration in cell supernatants over the course of the assay. In Fig. 4B the capture assay is compared to cell supernatant levels of IgM. There is correlation between on-cell capture and supernatant IgM concentration, with an R^2^ value for the line of best fit of 0.88 for the 1 hour assay and 0.93 for the 4 hour assay.

**Figure 4.**
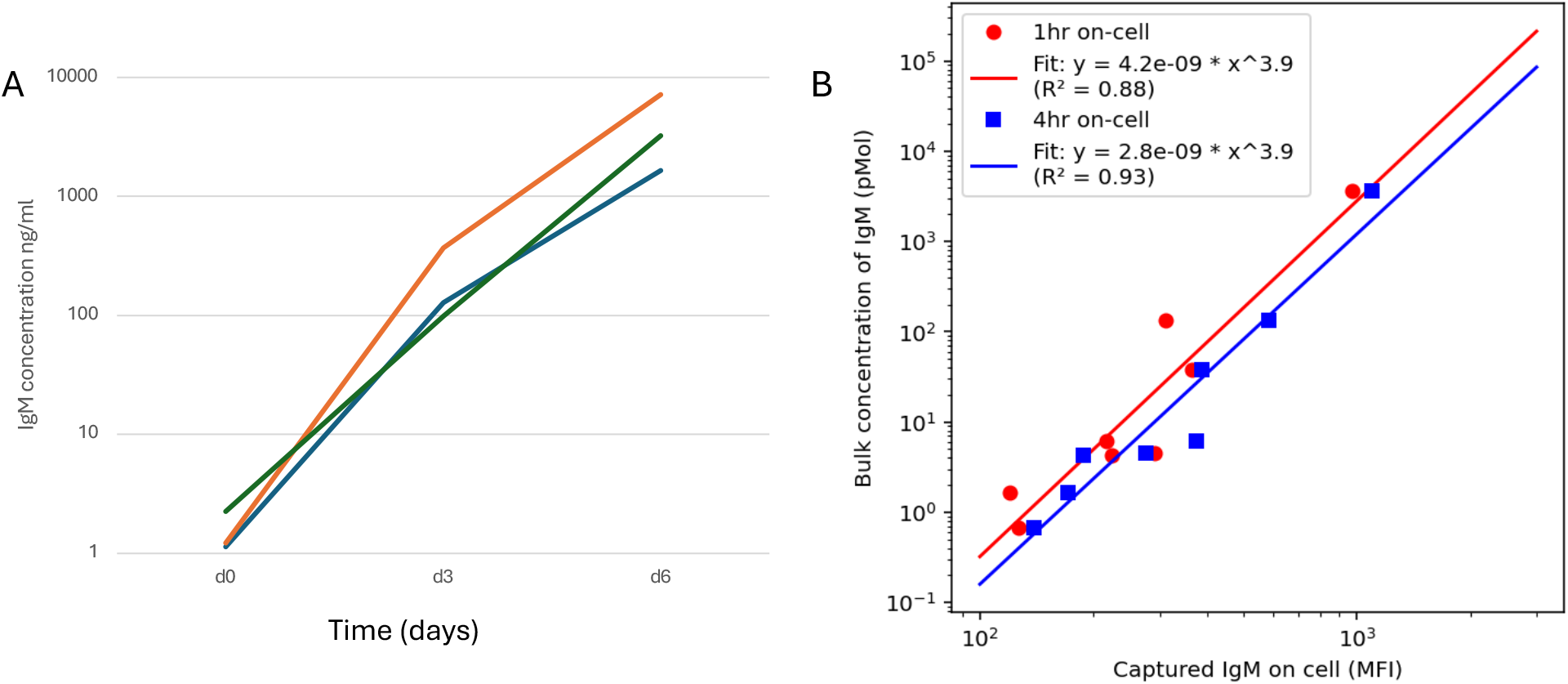
A bead ELISA assay demonstrates compares total Ig secretion in cell supernatant vs. the on-cell capture assay. A. B cells from 3 separate donors at differentiation time points as shown were spun out of the media and the collected media was tested for IgM levels using a bead ELISA assay. The lines show the individual donors. B. Concentration of IgM measured in cell supernatants was compared to captured IgM on cell (mean fluorescence intensity).

## Discussion

This work introduces and validates a streamlined single-antibody secretion capture assay that builds on, yet simplifies, existing secretion-capture approaches. Traditional secretion assays for antibodies and other secreted proteins typically rely on paired reagents: one reagent tethered to the cell surface (or a particle) to capture secreted protein, and a second, differently specific reagent to detect captured material.

Examples include loaded nanoparticles, covalently coupled capture/detection antibody pairs and plate-based techniques such as ELISPOT or membrane-capture formats^3,14^. Our approach leverages a fundamental biological feature of B cells: the membrane-bound and secreted isoforms of a given immunoglobulin share the same large variable and constant regions and therefore retain common epitopes. By using a bivalent anti-Ig reagent bound to surface BCR, one Fab arm anchors to membrane Ig while the other remains available to capture the secreted IgM molecule, which is then read out by a labelled anti-IgM secondary.

With our assay we have demonstrated that secretion can be measured on a cell-by-cell basis. Incubation on ice or with BFA as a control to inhibit secretion demonstrates that the detected signal is related to secretion, that increasing the time of assay also increases the level of secreted IgM detected, and that cells at different stages of differentiation secrete different levels of IgM. This demonstrates the workability of the assay for a live-cell secretion assay on primary human B cells.

Compared with existing secretion assays, this strategy offers several improvements. Firstly, it reduces reagent complexity and cost, by eliminating the need for antigenically distinct capture and detection antibodies or nanoparticle co-coupling. Secondly, because capture is effected directly at the native cell surface BCR, the assay preserves cell identity and spatial linkage between secretory phenotype and the originating cell; this could facilitate downstream single-cell phenotyping and sorting. Thirdly, the method is rapid, compatible with live cells and flow cytometry, and sensitive to dynamic changes in secretion (temperature dependence, stimulation time course and inhibition, e.g., by brefeldin A), enabling single-cell kinetic studies that many endpoint assays (e.g., ELISPOT, bulk supernatant ELISA) cannot resolve. Finally, because the assay does not perturb surface Ig levels over the timescale tested, signal changes reflect secretion rather than receptor internalization or turnover, addressing a common confounder in surface-capture approaches. Integration with cell sorting provides the opportunity to collect specific live cell populations of interest for downstream applications to allow for single-cell phenotyping^15,16^.

The assay has specific and broad utility in B cell biology. For IgM-producing B cells it provides a direct, single-cell readout of secretion during activation and differentiation, complementing transcriptomic and surface-phenotype analyses of plasmablast formation. It can be used to quantify how immunogens, adjuvants, or pharmacologic agents alter secretion kinetics. Since B cells are specifically adapted to the translation and then secretion of high volumes of antibodies, this assay offers a direct readout on the potential effect of treatments on the components directing these pathways. Similarly, it can also dissect mechanisms linking BCR engagement with secretory potential. Because the approach relies on shared epitopes between membrane and secreted forms, it should be extensible to other isotypes (IgG, IgA, IgE) provided suitable anti-isotype reagents that bind epitopes preserved in both isoforms are available; this opens the door to comparative studies of isotype switching and class-specific secretion dynamics. Practical applications include accelerated functional screening in antibody discovery (identifying high-secreting clones), monitoring vaccine responses or B cell-directed therapies in clinical studies, and coupling secretion phenotype with single-cell genomics or BCR sequencing to directly link antibody sequence, specificity and secretion capacity.

Future adaptations might incorporate isotype-specific secondary reagents for multiplexed detection, integration with microfluidic single-cell platforms for higher temporal resolution, or direct pairing with single-cell transcriptome and BCR sequencing to provide a comprehensive, multimodal map of B cell functional states^17,18^.

Here we capitalized on a native structural relationship between membrane and secreted immunoglobulins, which simplifies secretion measurement with single-antibody capture while preserving single-cell resolution and kinetic sensitivity. This assay therefore provides a practical, adaptable addition to the toolkit for studying B cell function and immunoglobulin biology, with translational and research applications across immunology and antibody discovery.

## Acknowledgements

The authors would like to thank Nihal Altan-Bonnet for sponsoring TW as a sabbatical researcher and for fruitful discussions on this project. We are grateful to Chris Chiu for support with flow cytometry. The work was supported by intramural NIH funds.

## Author Contributions

Conceptualization, methodology, investigation, writing (original draft and review & editing): TW and GAB; funding acquisition: GAB

## References

1. Nutt, S.L., Hodgkin, P.D., Tarlinton, D.M., and Corcoran, L.M. (2015). The generation of antibody-secreting plasma cells. Nat. Rev. Immunol. 15, 160–171. 10.1038/nri3795.

2. Boonyaratanakornkit, J., and Taylor, J.J. (2019). Techniques to study antigen-specific B cell responses. Front. Immunol. 10, 1694. 10.3389/fimmu.2019.01694.

3. Broketa, M., and Bruhns, P. (2021). Single-cell technologies for the study of antibody-secreting cells. Front. Immunol. 12, 821729. 10.3389/fimmu.2021.821729.

4. Brosterhus, H., Brings, S., Leyendeckers, H., Manz, R.A., Miltenyi, S., Radbruch, A., Assenmacher, M., and Schmitz, J. (1999). Enrichment and detection of live antigen-specific CD4+ and CD8+ T cells based on cytokine secretion. Eur. J. Immunol. 29, 4053–4059. 10.1002/(SICI)1521-4141(199912)29:12<4053::AID-IMMU4053>3.0.CO;2-C.

5. Fitzgerald, W., and Grivel, J.-C. (2013). A universal nanoparticle cell secretion capture assay. Cytometry A 83, 205–211. 10.1002/cyto.a.22199.

6. Trautmann, L. (2013). Beyond surface markers with a universal cell secretion assay. Cytometry A 83, 177–178. 10.1002/cyto.a.22200.

7. Clargo, A.M., Hudson, A.R., Ndlovu, W., Wootton, R.J., Cremin, L.A., O’Dowd, V.L., Nowosad, C.R., Starkie, D.O., Shaw, S.P., Compson, J.E., et al. (2014). The rapid generation of recombinant functional monoclonal antibodies from individual, antigen-specific bone marrow-derived plasma cells isolated using a novel fluorescence-based method. MAbs 6, 143–159. 10.4161/mabs.27044.

8. Pinder, C.L., Kratochvil, S., Cizmeci, D., Muir, L., Guo, Y., Shattock, R.J., and McKay, P.F. (2017). Isolation and characterization of antigen-specific plasmablasts using a novel flow cytometry-based Ig capture assay. J. Immunol. 199, 4180–4188. 10.4049/jimmunol.1701253.

9. de Rutte, J., Dimatteo, R., Archang, M.M., van Zee, M., Koo, D., Lee, S., Sharrow, A.C., Krohl, P.J., Mellody, M., Zhu, S., et al. (2022). Suspendable hydrogel nanovials for massively parallel single-cell functional analysis and sorting. ACS Nano 16, 7242–7257. 10.1021/acsnano.1c11420.

10. Kirk, S.J., Cliff, J.M., Thomas, J.A., and Ward, T.H. (2010). Biogenesis of secretory organelles during B cell differentiation. J. Leukoc. Biol. 87, 245–255. 10.1189/jlb.1208774.

11. Achar, S.R., Bourassa, F.X.P., Rademaker, T.J., Lee, A., Kondo, T., Salazar-Cavazos, E., Davies, J.S., Taylor, N., François, P., and Altan-Bonnet, G. (2022). Universal antigen encoding of T cell activation from high-dimensional cytokine dynamics. Science 376, 880–884. 10.1126/science.abl5311.

12. Cushley, W., Coupar, B.E.H., Mickelson, C.A., and Williamson, A.R. (1982). A common mechanism for the synthesis of membrane and secreted immunoglobulin **alpha, γ and µ chains. Nature 298, 77–79. 10.1038/298077a0.

13. Benjamin, D., Magrath, I.T., Maguire, R., Janus, C., Todd, H.D., and Parsons, R.G. (1982). Immunoglobulin secretion by cell lines derived from African and American undifferentiated lymphomas of Burkitt’s and non-Burkitt’s type. J. Immunol. 129, 1336–1342.

14. Bucheli, O.T.M., Sigvaldadóttir, I., and Eyer, K. (2021). Measuring single-cell protein secretion in immunology: Technologies, advances, and applications. Eur. J. Immunol. 51, 1334–1347. 10.1002/eji.202048976.

15. Saxena, A., Dagur, P.K., Desai, A., and McCoy, J.P., Jr. (2018). Ultrasensitive quantification of cytokine proteins in single lymphocytes from human blood following ex-vivo stimulation. Front. Immunol. 9, 2462. 10.3389/fimmu.2018.02462.

16. Siris, S., Gladstone, C.A., Guo, Y., Patel, R., Pinder, C.L., Shattock, R.J., McKay, P.F., Langford, P.R., and Bidmos, F.A. (2023). Increasing human monoclonal antibody cloning efficiency with a whole-cell modified immunoglobulin-capture assay (mICA). Front. Immunol. 14, 1184510. 10.3389/fimmu.2023.1184510.

17. Miwa, H., Dimatteo, R., de Rutte, J., Ghosh, R., and Di Carlo, D. (2022). Single-cell sorting based on secreted products for functionally defined cell therapies. Microsyst. Nanoeng. 8, 84. 10.1038/s41378-022-00422-x.

18. Heumos, L., Schaar, A.C., Lance, C., Litinetskaya, A., Drost, F., Zappia, L., Lücken, M.D., Strobl, D.C., Henao, J., Curion, F., et al. (2023). Best practices for single-cell analysis across modalities. Nat. Rev. Genet. 24, 550–572. 10.1038/s41576-023-00586-w.

